# HistoSweep enables cellular-resolution tissue quality control for gigapixel images in digital pathology and spatial omics

**DOI:** 10.64898/2026.01.30.702675

**Authors:** Amelia Schroeder, Xiaokang Yu, Wei Li, Musu Yuan, Liran Mao, Jiyuan Yang, Nadja Sachs, Bernhard Dumoulin, George X. Xu, Xunda Luo, Alexander Huang, Katalin Susztak, Tae Hyun Hwang, Humam Kadara, Lars Maegdefessel, Jiyang Yu, Mingyao Li

## Abstract

High-resolution histology images are indispensable for pathology and increasingly serve as the structural backbone for spatial omics. Yet whole-slide images (WSIs) frequently contain artifacts, acellular voids, and background regions that, when included in computational workflows, introduce noise, degrade model accuracy, and compromise biological interpretation. Existing tools provide only coarse foreground-background separation, leaving a gap in fine-grained quality control (QC). Here we present HistoSweep, a scalable framework that generates morphology-aware tissue masks at cellular resolution. By integrating density filtering, texture descriptors, and adaptive thresholding, HistoSweep systematically removes non-informative tissue regions while preserving biologically meaningful microstructures. It processes billion-pixel WSIs in minutes on standard CPUs, requiring no GPU acceleration, and is deployable across research and clinical settings. Across 25 WSIs spanning distinct tissues, disease states, and spatial omics platforms, HistoSweep consistently outperformed existing methods. It enhanced visualization and segmentation, improved virtual cell type predictions, and safeguarded spatial transcriptomics integrity by detecting transcript leakage and transcript-histology misalignment. By enabling fine-grained, scalable QC, HistoSweep provides a foundational preprocessing step for reliable and reproducible digital pathology and spatial omics analyses.

## Introduction

Histology slides remain a cornerstone of biomedical research and clinical diagnostics, providing high-resolution insights into tissue organization and disease states. Hematoxylin and eosin (H&E) staining, in particular, has been the standard for over a century^1, 2^, providing nuclei-specific contrast together with stromal and cytoplasmic features that are essential for diagnosis, disease characterization, and research applications^3-6^. The advent of whole-slide imaging has transformed these slides into gigapixel-scale digital files^6-8^, enabling remote review, quantitative analysis, and integration with spatial omics data. Modern scanners can digitize hundreds of slides daily, often generating files exceeding one gigabyte each, making H&E the most abundant and accessible source of high-resolution tissue information^9^.

With the explosive growth of digital pathology, automated computational pipelines have been developed to extract histology-derived features that support a wide variety of downstream tasks, including tissue segmentation, spatial omics integration^10-14^, and inference of clinical properties such as grading, prognosis, and risk stratification^15-19^. These features span from traditional morphometrics, such as nuclear shape, density, and texture, to high-dimensional embeddings learned from deep learning models. More recently, pathology image foundation models such as HIPT^20^, Virchow^21^, UNI^22^, and Prov-GigaPath^23^, trained on hundreds of thousands of slides and over a billion image tiles, have produced transferable histology representations with broad applicability. However, the quality and biological relevance of these features are fundamentally constrained by the underling WSIs, which often contain large regions of non-biological material or artifacts. When such regions are included in downstream analyses such as clustering, cell-type classification, spatial annotation, and virtual spatial transcriptomics (ST), they can introduce noise, degrade predictive accuracy, and compromise biological interpretation.

Despite these challenges, most histology-based and spatial omics workflows still rely on coarse “standard masks” that only distinguish foreground tissue from background. These masks are typically generated using fast, simple, and contrast-based techniques, which are highly sensitive but prone to errors. Vision Transformer (ViT)-based methods such as those in iStar’s masking module^24^ improve robustness but remain limited to coarse delineations that fail to distinguish biologically meaningful structures. More advanced QC tools, such as EntropyMasker^25^ and GrandQC^26^, can detect broad tissue regions and major artifacts, but cannot reliably filter finer structures such as tissue tears, holes, acellular voids, or fluid-filled regions lacking cellular information. In addition, these methods often require GPU resources and extended runtimes, greatly limiting their scalability for large-scale studies.

These limitations highlight the need for scalable, fine-grained approaches that move beyond coarse foreground-background segmentation to generate biologically meaningful tissue maps. To address this gap, we present HistoSweep, a reference-free framework for efficient image filtering and QC at cellular resolution. HistoSweep integrates patch-level color statistics, texture descriptors, and adaptive thresholding to generate fine-grained, morphology-aware tissue masks directly from WSIs. Designed with scalability in mind, HistoSweep can process images containing up to three billion pixels in minutes on a standard CPU. By operating at user-defined patch resolutions (e.g., 16 × 16, 64 × 64, 224 × 224 pixels), HistoSweep aligns naturally with patch sizes used in pathology image foundation models^20-22^, enabling seamless integration with state-of-the-art histology embeddings.

HistoSweep supports a broad range of applications, including morphology-preserving image segmentation, improved visualization, transcript integrity and transcript-image alignment assessment in spatial transcriptomics, and H&E-based molecular profile prediction in broader spatial omics. By delivering biologically informed, cellular-resolution image filtering at scale, it elevates the quality of histological input and ensures more accurate, reproducible, and scalable spatial analyses for both discovery and translational applications in digital pathology and spatial omics.

## Results

### Overview of HistoSweep

HistoSweep integrates measures of spatial heterogeneity, texture analysis, and adaptive thresholding in a multi-step pipeline to automatically generate cellular-resolution tissue masks optimized for digital pathology and spatial omics applications (**Fig. 1**). The workflow begins with an H&E-stained WSI and proceeds through five major steps to produce a fine-grained tissue mask. In the first step, the input WSI is divided into a uniform grid of non-overlapping patches (16 × 16 pixels by default), referred to as superpixels. From each superpixel, HistoSweep computes intensity and variability statistics to construct an empirical joint distribution, where sparse, low-density regions, defined by the joint distribution of global-weighted intensity and local standard deviation across all patches (see **Methods**), are flagged as candidate non-informative or artifact-associated superpixels. In the second step, these candidate superpixels are subjected to density-based filtering to remove those that lack cellular or morphological content, thereby reducing the influence of acellular voids and gross artifacts. In the third step, the low-density superpixels identified in step two, are further characterized by combining color signatures with Gray Level Co-occurrence Matrix (GLCM)-based textural descriptors. These features are clustered using Gaussian mixture modeling (GMM) to separate structurally uniform, low-information regions from variable, biologically informative ones. In the fourth step, the remaining superpixels are further processed with adaptive Otsu’s thresholding, guided by a ratio metric that captures both local variation and intensity to ensure accurate delineation of cellular structures. Finally, in the fifth step, spatial connectivity analysis is applied to eliminate residual debris, background noise, and isolated small fragments, producing the final tissue mask.

**Fig. 1:**
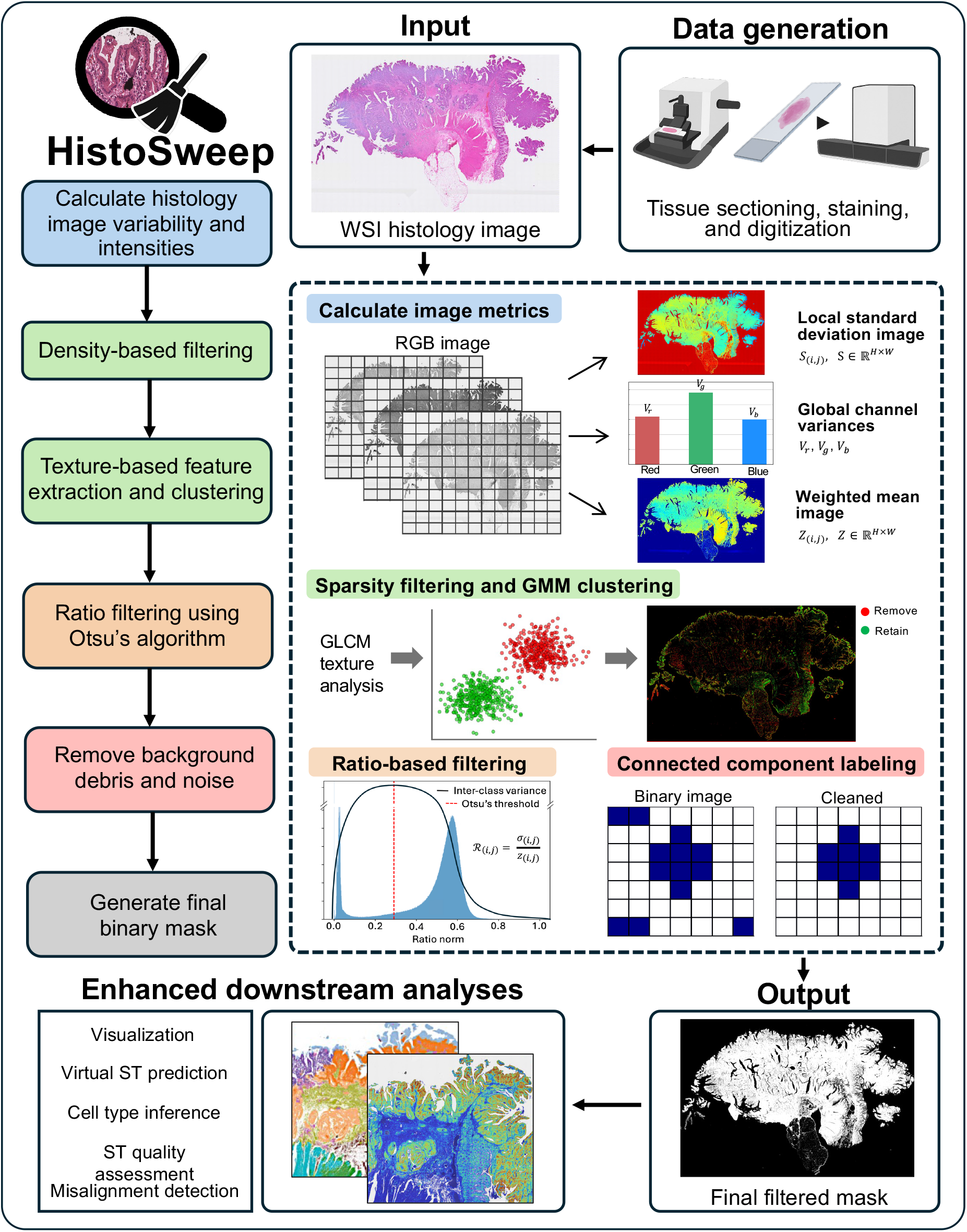
HistoSweep workflow. Overview of HistoSweep, a computationally efficient, fine-grained histology image filtering pipeline that computes local variability and intensity metrics to identify and remove artifacts, noise, and low-quality regions from histology images. The resulting binary mask enables enhanced downstream digital pathology tasks.

The outcome of this multi-step workflow is a fine-grained, morphology-aware tissue mask at cellular-resolution, selectively retaining biologically meaningful regions while excluding artifacts, voids, and other non-informative areas. HistoSweep’s stepwise design ensures robustness and scalability, delivering biologically grounded tissue masks for digital pathology and seamless integration with spatial omics data.

### Refining tissue masks through multi-stage filtering

Having introduced the overall workflow of HistoSweep, we next demonstrate why multi-stage filtering is necessary. WSIs often contain subtle, artifact-prone regions that global approaches fail to resolve, yet which can strongly influence downstream analyses. In this section, we illustrate why each refinement step, including density-based filtering, texture analysis, and adaptive thresholding, is required to systematically exclude low-content regions while preserving biologically informative structures.

The first challenge in WSI analysis arises from large areas of tissue that appear sparse in pixel intensity but may still contain a mixture of artifacts and biologically relevant compartments. To address this, HistoSweep begins with a density-based refinement workflow. In a kidney section with both healthy and type 2 diabetic (T2D) samples, where the H&E image was acquired after a CosMx experiment, an initial low-density filtering step was applied. This step identified superpixels located in sparse regions of the empirical joint distribution derived from global variance-weighted mean and local standard deviation RGB statistics (**Fig. 2a**). The resulting low-density candidates, visualized in blue (**Fig. 2b**), often corresponded to artifact-prone regions such as blurred areas, pen marks, or staining artifacts.

**Fig. 2:**
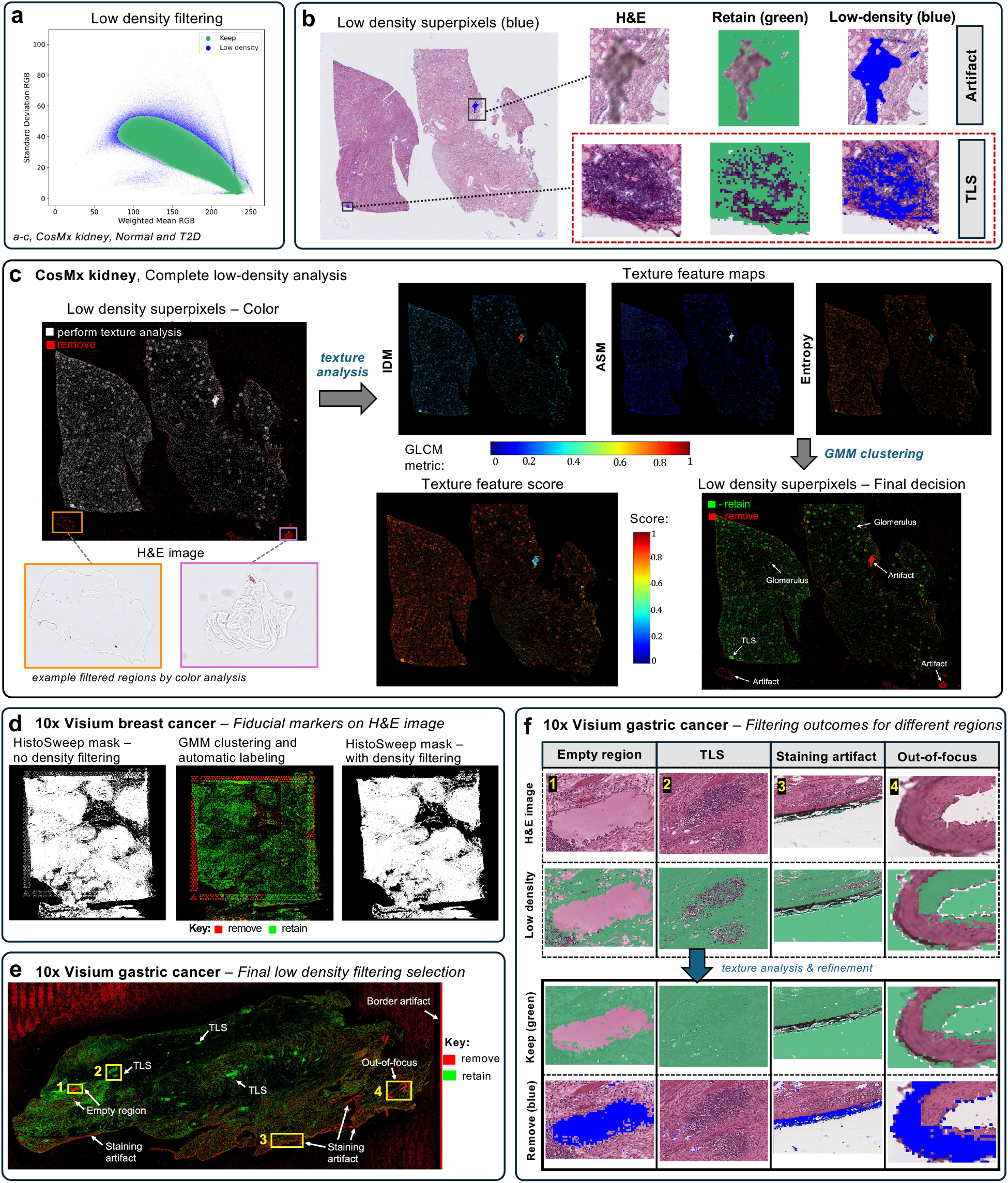
Multi-stage mask refinement in kidney, breast cancer, and gastric histology images. **a-c**, Analysis of multi-sample CosMx kidney H&E image. **a**, Scatter plot of the weighted mean versus local standard deviation metrics colored by low-density selection. **b**, Example of low-density regions including artifacts (top) and TLS (bottom). **c**, Color abnormality screen, computation of texture feature maps (IDM, ASM, and entropy), texture feature score, and final filtering decision for the set of low-density superpixels. **d**, Application of low-density filtering to remove fiducial grid artifacts in Visium breast cancer data. demonstrating robustness across tissue types and post-spatial omics imaging processing conditions. **e-f**, Gastric tumor post Xenium H&E image. **e**, Final filtering decision for the set of low-density superpixels. **f**, Filtering outcomes for different regions before and after texture feature analysis.

However, density filtering alone can sometimes flag biologically meaningful structures but visually outlying regions such as tertiary lymphoid structures (TLS), which contain densely packed, dark-staining nuclei with atypical intensity distributions. To prevent loss of such regions, HistoSweep incorporates a second refinement step based on color and texture features. Low-density superpixels are screened for coloration. As shown in **Fig. 2c** (CosMx kidney), gray-appearing imaging artifacts were removed during this step (red), while the remaining set of low-density superpixels (yellow) undergo GLCM-based texture analysis. Texture features, including inverse difference moment (IDM), angular second moment (ASM), and entropy, captured complementary aspects of spatial intensity variation. These features were clustered using a GMM, and a composite texture-feature score was computed to assign clusters to the decision groups: retain or remove, as described in **Methods**. As shown in **Fig. 2c**, this step successfully separated nuclei-rich structures such as TLS and glomeruli (green) from acellular voids and artifacts (red), producing a biologically meaningful decision map.

This approach generalized across tissues and platforms. In breast cancer tissue, the low-density filtering workflow successfully removed fiducial markers commonly found in 10x Visium datasets, while retaining surrounding epithelium (**Fig. 2d**). In gastric tissue, combined low-density and texture filtering eliminated empty regions, staining artifacts, out-of-focus areas, border artifacts, and background noise, while preserving TLS and other nuclei-rich compartments (**Fig. 2e,f**).

Even after density and texture-based filtering, residual low-information regions and background noise may remain, particularly at tissue edges or in regions with staining variability. To resolve this, HistoSweep applies a third refinement step based on adaptive ratio thresholding. This ratio metric captures the normalized relationship between local variation and intensity (see **Methods**), enabling effective removal of subtle background while retaining cellular structures. Representative distributions of the ratio metric and corresponding Otsu-derived thresholds across diverse tissues are shown in **Supplementary Fig. 1**.

Together, these results highlight the importance of sequential refinement steps for generating meaningful tissue masks and demonstrate how each stage of HistoSweep contributes to robust tissue filtering.

### Demonstrating robustness across tissues and spatial omics platforms

After establishing the rationale for multi-stage filtering, we next evaluated the robustness and reliability of HistoSweep across diverse biological and technical conditions. To this end, we evaluated its performance on 25 WSIs representing a wide spectrum of variation (**Fig. 3**; **Supplementary Table 1**). The datasets included human and mouse samples spanning 12 tissue types: colon, kidney, liver, stomach, breast, small intestine, artery, tonsil, lung, skin, bone marrow, and whole-body mouse sections, together with multiple disease contexts such as colorectal, breast, liver, gastric, and kidney cancers, melanoma, atherosclerosis, renal carcinoma, and T2D, as well as multiple healthy controls.

**Fig. 3:**
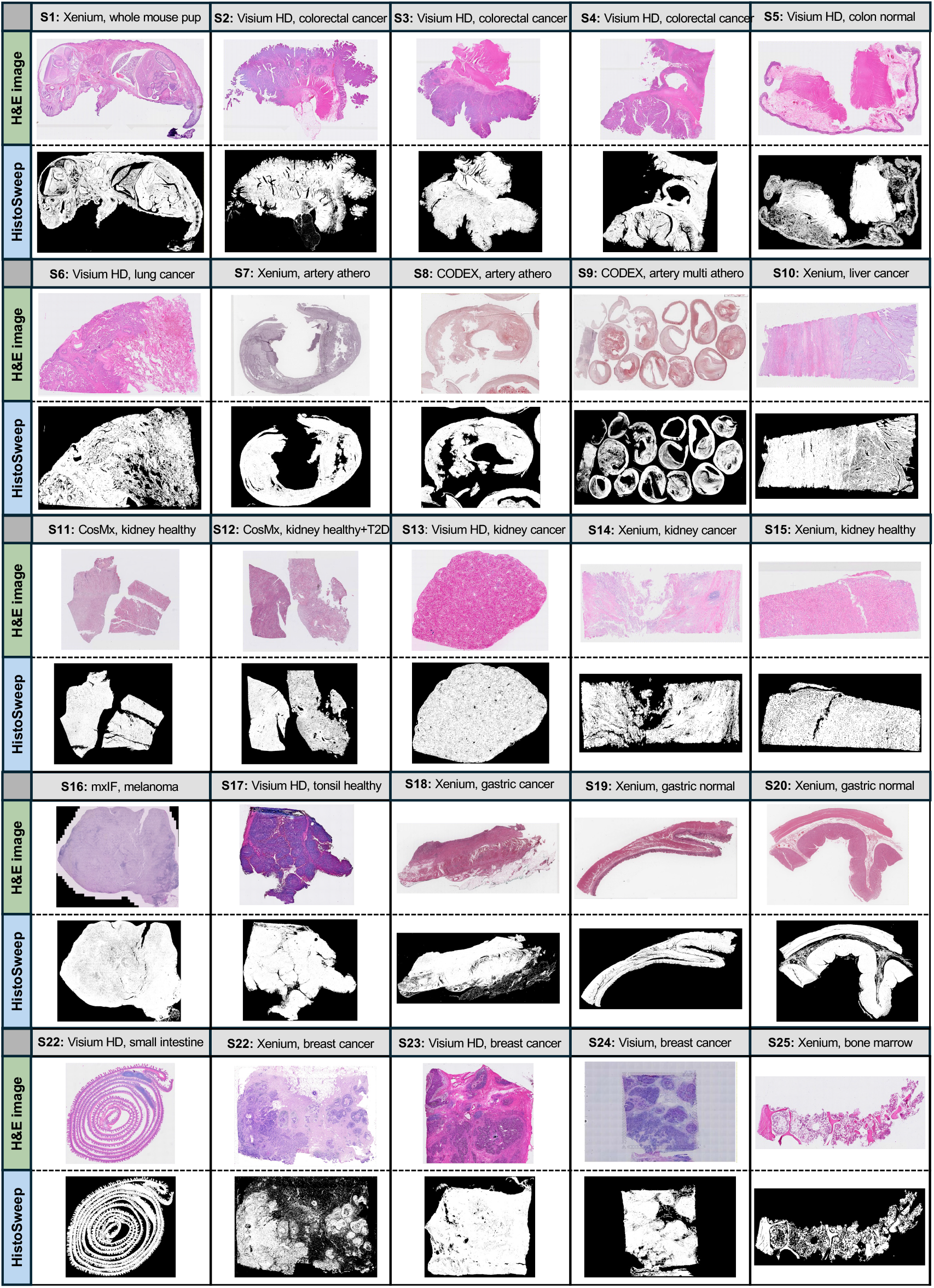
HistoSweep results across diverse tissue samples. HistoSweep results across all tissue sections analyzed in this study. Visualization of the H&E image and HistoSweep generated masks. The dataset covers 25 samples across a wide range of tissue types, disease states, staining concentrations, image sizes, and spatial omics platforms. Corresponding sample descriptions and metadata are provided in Supplementary Table 1.

To capture real-world variability, we analyzed images with markedly different H&E staining concentrations (**Fig. 3**), ranging from low concentration (e.g., S8-CODEX, atherosclerotic artery) and highly saturated (e.g., S17-Visium HD, tonsil). Both fresh frozen and FFPE preparations were included to assess robustness across preservation techniques. Image dimensions varied substantially, from 4,864 × 15,104 pixels (S14-Xenium, kidney cancer) to 73,728 × 40,704 pixels (S9-CODEX, multi-sample atherosclerotic artery), with some images exceeding three billion pixels (**Supplementary Fig. 2**). These analyses highlight HistoSweep’s scalability to ultra-large and high-resolution images.

From a technological standpoint, the analyzed datasets encompassed images generated either before or after spatial omics protocols, including 10x Genomics Xenium, 10x Genomics Visium HD, NanoString CosMx, and Akoya CODEX. Images obtained after spatial omics profiling often exhibited evidence of tissue degradation, morphological artifacts, or diminished image quality. Despite these challenges, HistoSweep consistently generated meaningful and accurate masks, highlighting its resilience to pre-analytical variation and imaging artifacts.

Together, this broad and heterogeneous benchmark demonstrates that HistoSweep is robust to variation in tissue type, disease state, image size, staining quality, preservation method, and spatial omics platform, enabling consistent tissue filtering and QC across a broad range of histology datasets.

### Comparison with existing tissue masking methods

To compare HistoSweep with existing approaches, we compared it with the recently published H&E image QC tools: GrandQC^26^ and the ViT-based masking tool from iStar^24^, which generate the commonly used foreground-background “standard” mask. Five diverse WSIs were analyzed (**Fig. 4**), spanning a broad range of tissue types, staining conditions, and spatial omics protocols. This range enabled a comprehensive evaluation of each method’s ability to retain biologically meaningful tissue while excluding irrelevant or low-quality regions. Notably, GrandQC could not process several of these gigapixel images without code modification, and it requires GPU-enabled execution, highlighting its scalability limitations.

**Fig. 4:**
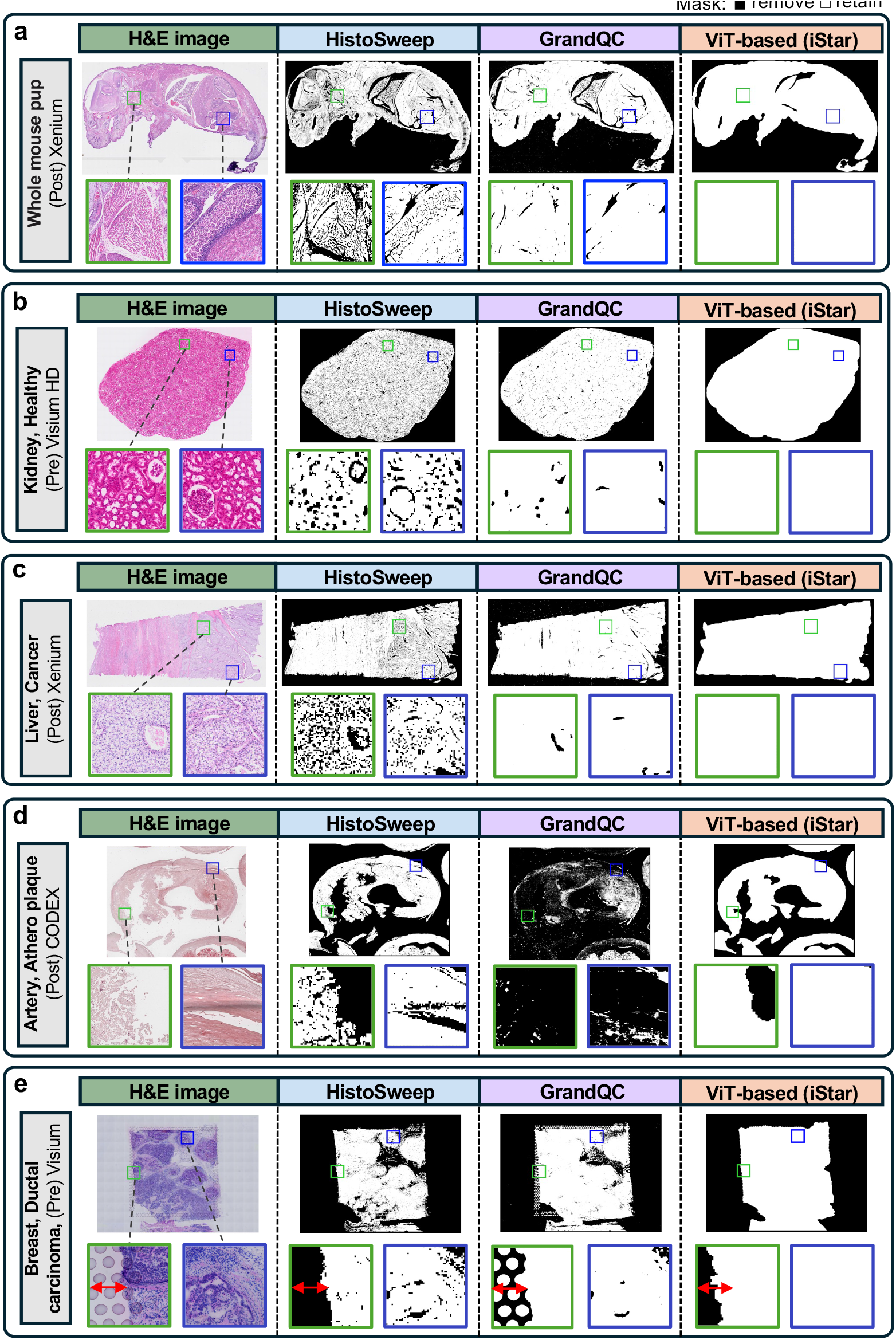
Comparative evaluation of HistoSweep, GrandQC, and ViT-based standard masking across diverse tissue types. Representative H&E images (left) alongside corresponding binary masks (black = filtered, white = retained) generated by HistoSweep, GrandQC, and a ViT-based iStar mask for **(a)** whole mouse pup (10x Xenium, post-ST), **(b)** healthy kidney (10x Visium HD, pre-ST), and **(c)** liver cancer (10x Xenium, post-ST). Insets indicate example regions used to assess tissue structures and masking output. **(d)** atherosclerotic artery (CODEX, post), **(e)** ductal carcinoma breast cancer (10x Visium, pre-ST). Red arrow indicates the start of the tissue boundary compared to the fiducial markers. Insets indicate example regions used to assess tissue structures and masking output.

In the case of the whole mouse pup, post-Xenium H&E image (**Fig. 4a**), which represents a large and anatomically complex sample containing structures spanning the head, thorax, abdomen, limbs, and connective tissue, HistoSweep captured detailed anatomical boundaries and preserved internal cell-rich regions while excluding extensive non-informative spaces (**Supplementary Fig. 3**). In contrast, GrandQC retained large internal spaces devoid of cellular architecture, reducing filtering precision in areas such as the cranial and abdominal cavities, and failed to filter a prominent staining artifact in the bottom right corner. For all samples analyzed, the ViT-based mask produced an over-smoothed outline of the tissue, failing to resolve fine anatomical boundaries or exclude non-cellular internal spaces (**Fig. 4**).

A similar trend was observed in the healthy kidney sample (**Fig. 4b**). Here, HistoSweep successfully preserved fine-grained internal structures, including glomeruli, corpuscle, and tubules, while removing the surrounding empty regions and acellular fluid-filled space. In contrast, GrandQC tended to over-retain low-information internal voids and isolated luminal spaces and failed to capture finer architectural variations within the renal cortex and renal medulla.

The comparison in the Xenium liver cancer sample (**Fig. 4c**) further illustrates these differences. HistoSweep selectively retained regions of tumor-infiltrated liver parenchyma, where hepatocyte plates appear compressed and irregular, consistent with features of hepatocellular carcinoma (HCC)^27^. The right side of the tissue shows disorganized cellular architecture with widened intercellular spaces and distorted plates, suggesting tissue remodeling or early fibrosis. When investigated, these regions include morphologically atypical, *GPC3*-high tumor areas. HistoSweep appropriately filtered intervening low-content regions, such as necrotic zones or non-informative voids, enhancing structural continuity and contrast, which is critical for downstream interpretation of cellular patterns in the tumor microenvironment. In contrast, the right side of the tissue, retained more extensively by HistoSweep, displays preserved hepatic architecture and elevated *CYP3A4* expression, a marker of normal hepatocyte function. GrandQC, however, retained broad regions spanning both tumor and normal areas with limited structural discrimination.

Additional examples further highlight HistoSweep’s advantages. In the atherosclerotic artery sample (**Fig. 4d**), HistoSweep captured necrotic tissue structures and preserved vessel morphology while excluding background, holes, and imaging artifacts. For instance, in the blue-box region, a prominent out-of-focus blur line is successfully removed by HistoSweep. In contrast, biologically relevant regions were largely over-filtered by GrandQC. Similarly, in the breast ductal carcinoma sample (**Fig. 4e**), HistoSweep successfully removed the common fiducial grid patterns present in 10x Visium slides and selectively retained morphologically relevant tumor epithelium. In contrast, GrandQC and the “standard” mask failed to separate the fiducial grid patterns artifacts from tissue. Overall, these examples collectively demonstrate that HistoSweep consistently achieves finer grained and morphologically aware filtering compared to existing tools.

### Enhancing tissue visualization

In digital pathology and spatial omics, high-resolution tissue visualization is essential not only for aesthetic clarity but also for accurate interpretation of spatial features and morphological patterns, including interpreting predicted regions, inspecting model outputs, or validating annotations. Poor-quality areas such as tissue holes, folds, or staining artifacts can visually obscure important structures and hinder downstream review. This becomes especially problematic when downstream analyses rely on image-derived features or when spatial omics data are low-resolution or noisy.

The default ViT-based iStar mask produces a type of tissue mask that closely resembles the commonly used standard mask; however, as shown previously, it tends to over-smooth and obscure fine structures by considering only foreground versus background. To address this limitation, HistoSweep enables fine-grained filtering that produces detailed tissue masks that enhance visualization while preserving biologically meaningful content. **Fig. 5** demonstrates the substantial improvement in tissue visualization achieved when using HistoSweep compared to a standard mask, across six diverse samples, including both healthy and diseased tissues from mouse and human specimens. Notably, in the small intestine, HistoSweep preserved fine-grained structures such as epithelial folds of intestinal villi while also removing artifacts and gaps introduced by the Swiss roll preparation technique outlined in the protocol (S21)^28^. It also captured trabecular patterns in bone marrow (S25), and bronchial structures in lung cancer (S6) samples. In the whole-mouse pup tissue (S1), HistoSweep enhanced the spatial delineation of developing organs, while in non-diseased colon and colorectal cancer tissues (S5, S2), it maintained the architectural integrity of colonic crypts structures and mucosal layers.

**Fig. 5:**
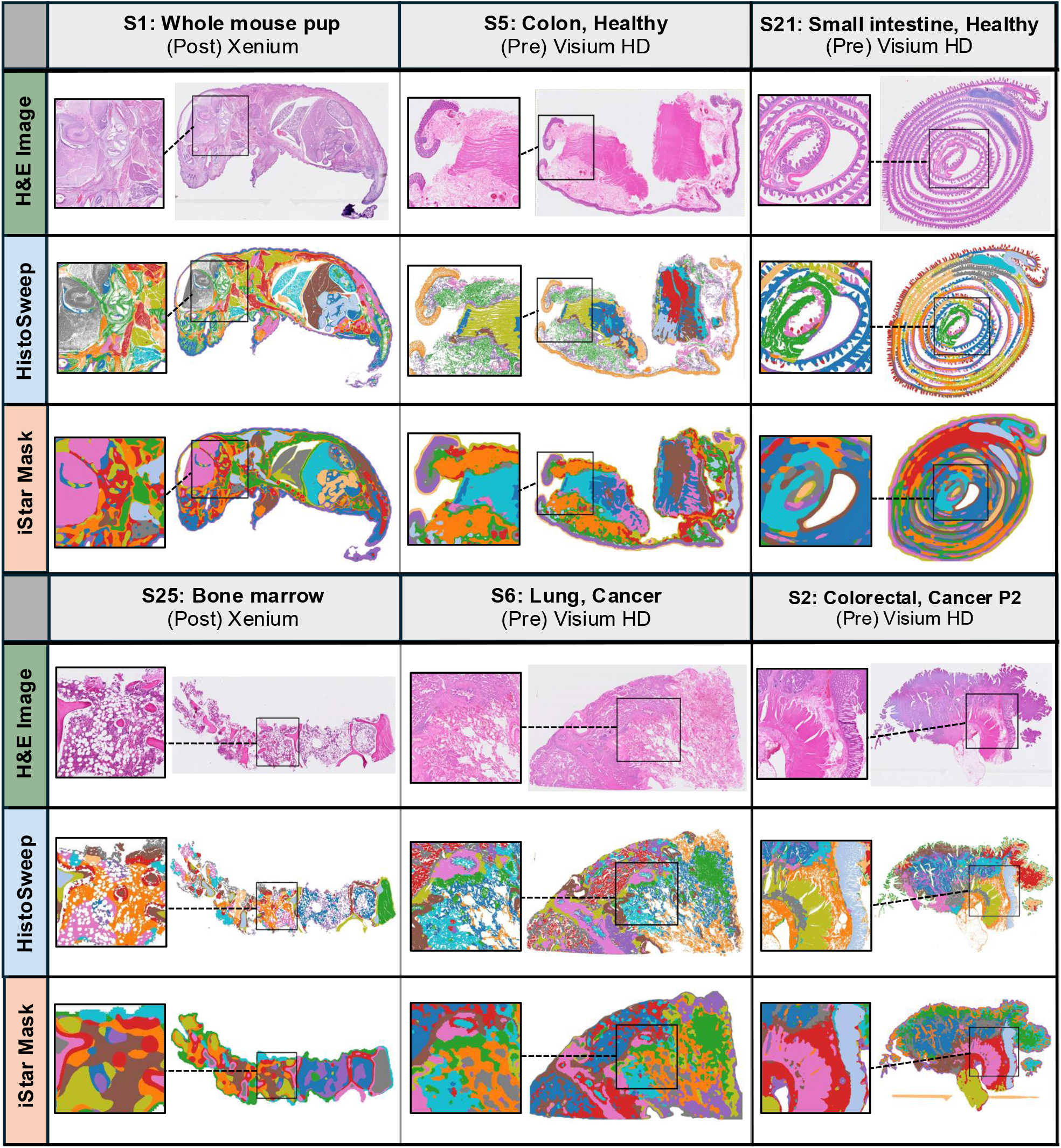
Visualization of tissue architecture segmentation. Comparison of tissue clustering results and visualizations using a standard-type mask compared to HistoSweep mask across six samples. Histology-derived HIPT features were extracted for each sample, with the same feature set used for both masks. These masks were applied independently before clustering. Insets show corresponding H&E images and resulting clustering maps.

These results highlight that HistoSweep is not limited to the removal of background or staining artifacts but also preserves morphologically and biologically informative structures that are critical for contextualizing spatial omics results and interpreting tissue organization at scale. This underscores the importance of accurate and detailed tissue masking as a foundational preprocessing step in spatial analyses.

### Reducing transcript signal leakage in ST

HistoSweep not only enhances visualizations but also helps assess and improve ST data integrity by pinpointing fine-grained regions like tissue tears, holes, and acellular voids that should not exhibit high transcript signals. Spatial omics technologies, including grid-based ST platforms like Visium HD, can suffer from signal leakage, which occurs when transcripts or background signals are mistakenly assigned to incorrect spatial locations. This misassignment can lead to inflated expression values in non-tissue regions and distorted spatial expression profiles. Potential leakage can stem from multiple technical and biological factors, including ambient RNA contamination, permeabilization inconsistencies that allow transcripts to diffuse across boundaries, tissue misregistration, capture resolution, or imprecise capture area boundaries ^29-31^. Notably, 10x Genomics Visium HD technical note recommends deep sequencing (∼275 million reads per capture area for FFPE samples) to saturate transcript detection, highlighting the importance of accurate capture despite the risk of contamination or misassignment.

**Fig. 6** demonstrates the utility of HistoSweep in identifying high-UMI regions that lack histological support in healthy kidney and lung cancer tissues. This analysis focused on “bins under tissue” as defined by the 10x Visium HD software. In the healthy kidney sample (**Fig. 6a**, left column), the H&E image highlights two regions (Box 1 and Box 2) that include Bowman’s space, a fluid-filled, acellular compartment surrounding the glomerulus, where little to no transcript expression is expected in healthy tissue^32^. However, the Visium HD data show elevated UMI counts in these regions, despite the absence of supporting cellular structures. By contrast, adjacent glomeruli, which contain dense and transcriptionally active cells, represent the biologically expected sites of high expression. The close proximity of Bowman’s space to these densely transcribed glomeruli suggests that the observed high UMI signal in this acellular region may result from signal spillover or diffusion. Hollow regions within luminal or tubular spaces that lack nuclei were also effectively filtered, while surrounding tubular epithelial cells with intact nuclei and transcriptional activity were preserved. This spatial selectivity, which excludes the acellular interior while retaining biologically relevant lining, offers a more accurate alternative to approaches that indiscriminately mask entire tubules.

**Fig. 6:**
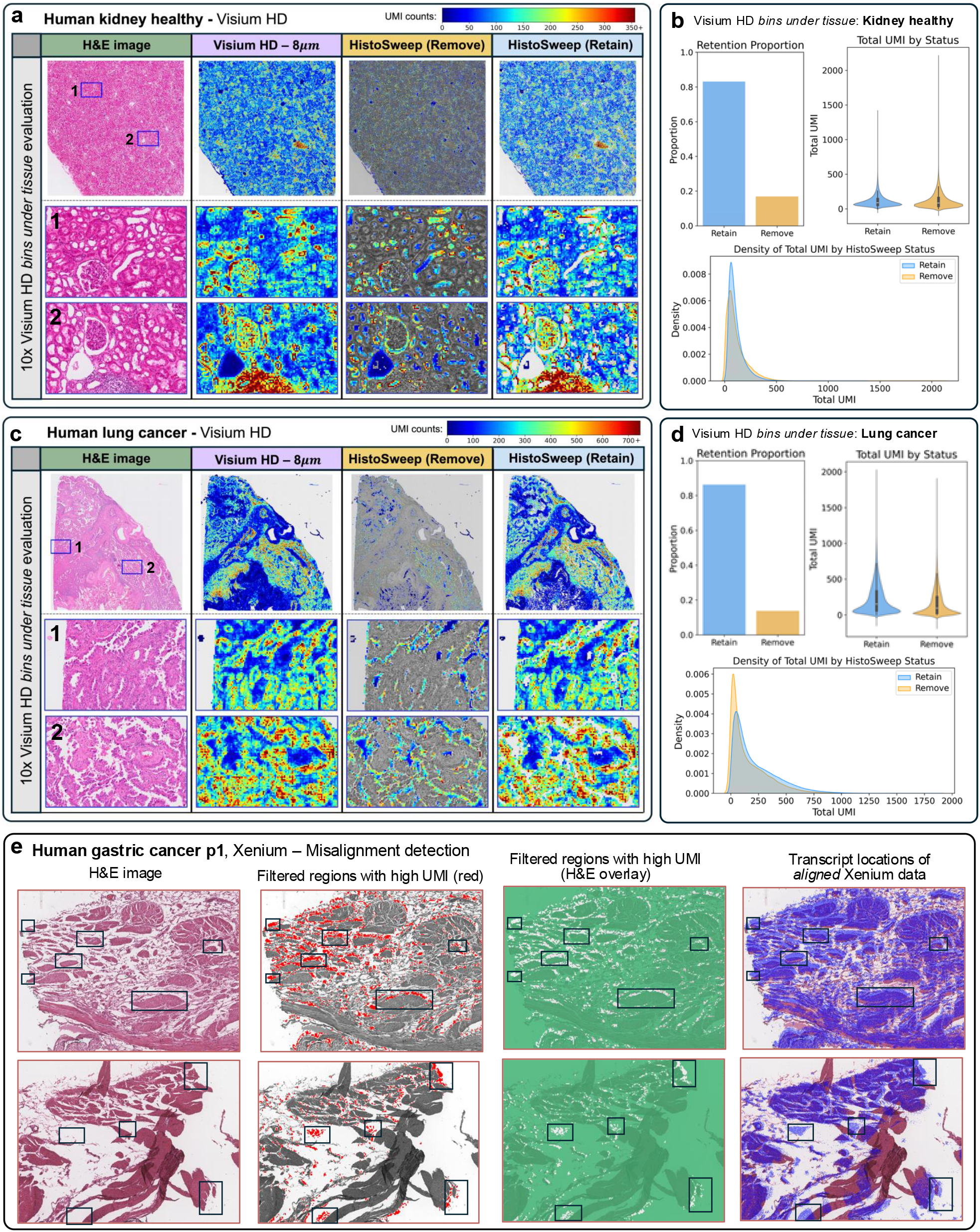
HistoSweep enables detection of signal leakage in Visium HD kidney and lung cancer data and misaligned high-UMI regions in a Xenium gastric cancer sample. Analysis using only “bins under tissue” determined by Visium HD software. **a, c**, H&E images, Visium HD total UMI counts (8□µm bins), and HistoSweep removal and retention maps for healthy human kidney (a) and lung cancer (c) samples. Insets show magnified regions highlighting local histology and corresponding transcript counts. **b, d**, Proportion of bins retained and removed by HistoSweep (left), violin plots showing total UMI by HistoSweep status (right), and density distributions of total UMI counts across retained and removed bins (bottom) for kidney (b) and lung (d) samples. **e**, Total UMI vs. ratio statistic plot highlighting superpixels with high UMI and low ratio (left) and calculated HURR = 6.33%, along with their spatial distribution across the tissue section (right). **f**, H&E image and corresponding overlays showing spatial locations of HistoSweep filtered high-UMI superpixels (red), with evidence of misalignment between transcript locations and histological structures.

A similar pattern was observed in the lung cancer sample (**Fig. 6c**), where sharply demarcated white areas in Panels 1 and 2 corresponded to the empty lumina of pulmonary alveoli. These areas appear white on H&E sections due to the loss of air and fluid during tissue processing. Tumor cells proliferate along the preserved alveolar septa, leaving the air spaces intact and often slightly expanded. Because these cavities lack resident cells, barcoded bins over them are expected to have low UMI and gene counts. However, Visium HD data revealed elevated UMI counts in these non-cellular spaces, likely reflecting ambient RNA contamination or accidental overlay of thin tissue folds and should be flagged during QC.

The distributions of UMI counts across Visium HD’s “bins under tissue” further demonstrate that superpixels filtered out by HistoSweep, despite corresponding to regions with minimal or no discernible cellular material, exhibit expression profiles similar to those of retained regions in both samples (**Fig. 6b,d**). This suggests notable signal leakage or contamination in the raw Visium HD measurements. In the kidney sample, many false-positive expression signals were observed across luminal and acellular fluid-filled areas, particularly near densely cellular regions, leading to high UMI counts in the filtered regions (**Fig. 6b**). The lung cancer sample showed a slightly improved pattern, with false-positive signals more spatially localized, resulting in filtered superpixels exhibiting slightly lower UMI counts and a stronger peak near zero in the distribution (**Fig. 6d**). Nonetheless, residual false positives were still apparent in some non-tissue regions.

We also investigated a third Visium HD dataset from a colorectal cancer sample, which exhibited substantially less evidence of transcript leakage or false-positive signals (**Supplementary Fig. 4**). In this case, the violin plot revealed a clear separation in total UMI counts between bins retained and removed by HistoSweep, with filtered bins showing significantly lower transcript abundance, as biologically expected. This is further supported by the density distribution (**Supplementary Fig. 4b)**, where most filtered bins have UMI counts near zero. The few remaining false positives were primarily located along tissue borders, suggesting minor edge effects.

Lastly, we compared bins removed by HistoSweep with those labeled as “not under tissue” by the Visium HD software, treating these as true negatives. Across all samples, HistoSweep achieved a true negative rate above 0.99, indicating strong agreement in identifying background regions (**Supplementary Fig. 5a,c,e**). When applied to all bins in the image, HistoSweep produced a UMI distribution more consistent with biological expectations: removed bins were predominantly low or near zero in UMI, while retained bins exhibited higher transcript levels (**Supplementary Fig. 5b,d,f**).

### Identifying transcript-histology misalignment

We also explored the relationship between total UMI counts and the HistoSweep ratio metric in three single-cell ST datasets generated using the 10x Xenium platform. We observed a strong correspondence between the ratio metric and total UMI, particularly in areas of high density (**Supplementary Fig. 6**). However, in the gastric cancer sample, we detected a notable subset of superpixels with high UMI that fell below the adaptive filtering threshold, placing them in the low-ratio, high-UMI region (**Supplementary Fig. 7a**, red zone). This group represented 6.33% of all high-UMI superpixels (i.e. high UMI removal rate, HURR), indicating a nontrivial removal rate, and raising the possibility of erroneously filtering biologically relevant signal. To assess these regions further, we examined their spatial context within the tissue and found that the filtered superpixels in this category were located in areas with visible holes, tissue tears, or regions devoid of tissue, consistent with structural artifacts, and confirming correct filtering by HistoSweep. Upon closer inspection, we discovered that the elevated UMI levels in these regions arose from misalignment between the transcript data and the histology image (**Fig. 6e**). These misaligned superpixels were often offset from dense tissue structures and histological boundaries, or located in areas where tissue may have detached from the slide.

**Fig. 7:**
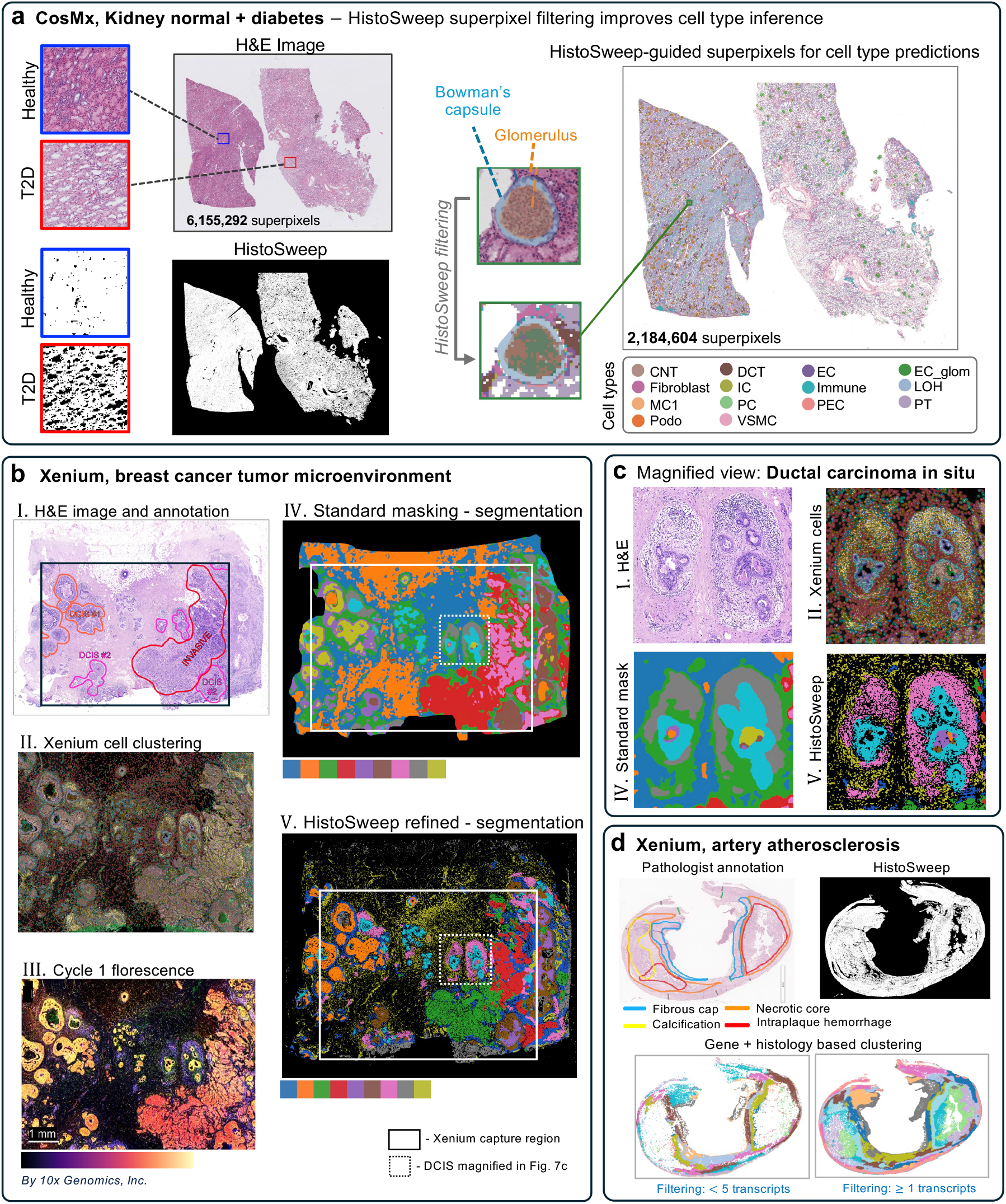
HistoSweep-enhanced downstream spatial analysis across multiple platforms and tissue types. **a**, H&E images, HistoSweep masks, and S2Omics cell type predictions guided by HistoSweep superpixel selection for CosMx kidney tissue from healthy and T2D donors. **b**, Pathologist annotation, HistoSweep tissue mask, transcript filtering, and gene□+□histology-based clustering in a Xenium human atherosclerotic artery sample. **c**, H&E image, Xenium cell clustering, fluorescence imaging, and segmentation outputs from standard imputed ST and HistoSweep-refined imputed ST predictions in a breast cancer sample. **d**, Magnified view of DCIS showing H&E, Xenium cell map, standard mask, and HistoSweep-refined segmentation.

Next, we applied the same analysis to two Xenium gastric normal samples to evaluate whether high-UMI superpixels below the ratio threshold also corresponded to poorly aligned tissue regions (**Supplementary Fig. 7**). In the first normal sample (P1), 3.12% of the high-UMI superpixels were filtered, with many of these regions localized near tissue border or epithelial edges (**Supplementary Fig. 7c**). Spatial inspection again revealed misalignment between the transcript data and the H&E image, including shifted transcript locations relative to epithelial cells. In contrast, the second normal sample (P2) had a significant lower HURR of 0.66%, and the transcript signals were well aligned with tissue architecture (**Supplementary Fig. 7d,e**). These findings suggest that when manual alignment is accurate, the HURR by HistoSweep is minimal.

Manual alignment remains an important preprocessing step when working with image-based spatial platforms such as Xenium. Given these observations, HistoSweep may serve as a complementary tool to automatically pinpoint regions of potential misalignment and provide a quantitative framework for evaluating alignment quality.

### Improving downstream spatial analyses

To further evaluate the utility of HistoSweep in enhancing downstream spatial analyses, we applied it to multiple tissue samples and spatial omics platforms, including CosMx, Xenium, and CODEX (**Fig. 7**). These examples highlight how HistoSweep filtering enables more accurate cell type inference, tissue segmentation, and structural delineation in spatial omics data.

In CosMx kidney tissue from both healthy and type 2 diabetic (T2D) individuals, HistoSweep was used to filter superpixels prior to cell type annotation (**Fig. 7a**). This substantially reduced the number of noisy, non-informative, or artifact-prone regions passed to the S2Omics^33^ analysis pipeline, which relies on H&E image derived features for virtual cell type prediction. In this dataset, the number of superpixels was reduced from 6,155,292 to 2,184,604, effectively excluding low-quality regions such as debris, tissue gaps, tears, and fluid-filled spaces (**Fig. 7a**). Downstream cell type predictions, guided by HistoSweep’s filtered superpixel selection, showed clearer spatial compartmentalization and improved alignment with kidney microstructures, including the glomerulus and the surrounding Bowman’s space and capsule. The refined superpixel set also enhanced fine-grained cell type assignment by preserving histologically relevant boundaries and reducing noise from acellular regions.

We next applied HistoSweep to a Xenium dataset of human atherosclerotic artery, where it generated a refined tissue mask that retained structurally complex regions such as necrotic core and intraplaque hemorrhage (**Fig. 7b**). These regions aligned with transcript presence filtering even in low-density areas, suggesting that HistoSweep may help validate biologically relevant structures, including the extent of necrosis or hemorrhage, while preserving spatial continuity across regions of interest with sparse but meaningful expression.

Lastly, in a Xenium dataset of human breast cancer tissue, HistoSweep improved segmentation of the tumor microenvironment, including regions of ductal carcinoma in situ (DCIS) (**Fig. 7c,d**). In both the standard and refined segmentations (panels IV and V), clusters were derived from imputed gene expression generated by the iStar^24^ software, where virtual ST prediction was performed on a benchmarking pseudo-Visium dataset. However, in the HistoSweep-refined version (panel V), the virtual gene expression was filtered using the HistoSweep mask prior to clustering. This resulted in cleaner input and more accurate segmentation (panel V), closely aligning closely with the true cellular regions (panels II and III) measured by ST, and preserving ductal boundaries and internal tumor structures. The improvement was especially apparent in the magnified DCIS region, where cell-level segmentation more closely matched true cellular contours and revealed internal heterogeneity that was captured by the standard mask (**Fig. 7d**).

## Discussion

In this study, we introduced HistoSweep, a scalable and modular framework for histology image QC that supports tissue quality assessment, enhances visualization, and enables robust downstream spatial analyses. By integrating color-based statistics, texture descriptors, and adaptive thresholding, HistoSweep generates fine-grained, morphology-aware masks that preserve biologically meaningful structures while removing irrelevant, low-content, and artifact-prone regions. The framework is computationally efficient, capable of processing a 3-billion-pixel image in just minutes on a CPU without requiring GPU resources, making it readily adaptable for both research and clinical workflows.

A major strength of HistoSweep is its broad generalizability across tissue types, disease states, imaging protocols, and staining quality. Across 25 diverse WSIs, including human and mouse tissues, pre- and post-spatial omics imaging, and challenging cases with varying staining quality, HistoSweep consistently produced accurate, morphology-preserving tissue masks (**Fig. 3**). Importantly, the results remained robust even under extreme conditions, including over-saturation, image degradation, and complex, unusual tissue architectures, enabling confident application to real-world digital pathology and spatial omics datasets. Compared to existing tissue masking tools such as GrandQC and standard masking such as the ViT-based models, HistoSweep demonstrated superior performance in retaining fine-grained tissue structures such as glomeruli, crypts, fibrotic boundaries, and tumor ductal structures, while filtering out artifacts, background noise, staining irregularities, and acellular areas.

A particularly notable application of HistoSweep is its utility in improving the interpretability and integrity of spatial omics data. In Visium HD data, HistoSweep identified histologically void regions with unexpectedly high transcript signal, revealing probable spillover or capture bias. Filtering such false-positive signals substantially reduced noise in ST fidelity of gene expression measurements. In imaging-based ST platforms like 10x Xenium, HistoSweep identified regions of high-UMI, low-ratio superpixels indicative of transcript-histology misalignment, providing a quantitative means to assess registration accuracy. These findings highlight the value of HistoSweep as a QC layer that safeguards the integrity of ST data.

Beyond spatial omics, HistoSweep significantly improved segmentation, tissue visualization, clustering, and cell type prediction in downstream image analysis tasks using histology derived features. Its computational efficiency is a key advantage: HistoSweep is capable of processing high-resolution WSIs, including multi-sample slides with over a billion pixels, in just minutes on a standard CPU, without the need for GPU acceleration or code modification. This scalability enables deployment in large-scale histology studies and clinical workflows where throughput is critical. In contrast, GrandQC requires GPU acceleration and was unable to run on large images without code modification.

In future work, HistoSweep could be extended to histological image types beyond H&E, such as immunohistochemistry (IHC), where its patch-based filtering could help identify and quantify marker-positive regions while excluding background or nonspecific signal. As illustrated in **Supplementary Fig. 8**, HistoSweep accurately localized marker-positive regions in IHC-stained human samples while preserving relevant structural detail, underscoring its adaptability beyond H&E. Broader applications may include multiplex imaging modalities, where morphology-aware filtering could support multi-marker interpretation and cell phenotyping. Additional future directions include leveraging learned representations from vision foundation models to enhance patch characterization and developing quantitative metrics to assess transcript-image alignment accuracy and transcript leakage based on HistoSweep outputs.

At the same time, HistoSweep has several limitations. While the framework produces fine-grained, morphology-aware masks, subtle biological features that lack clear histological cues may still be excluded. In addition, although computationally efficient on CPUs, scaling to extremely high-resolution or multi-channel datasets may require further optimization. Addressing these limitations will be key to extending HistoSweep’s applicability across wider range tissues, imaging modalities, and spatial omics platforms.

In summary, HistoSweep provides a computationally efficient and biologically informed solution for QC of H&E based WSIs. By enhancing the quality and utility of histological input, HistoSweep enables more accurate, reproducible, and scalable analyses in both digital pathology and spatial omics, advancing research and translational applications in tissue biology.

## Acknowledgments

M.L. was partly supported by the following NIH grants R01HG013185, R01LM014592, U19NS135582, R01HL171595, and U01CA294518. T.H.H. was partly supported by NIH grants R01CA276690 and U01CA294518, DOD grant CA190578, the Eric and Wendy Schmidt Foundation’s AI Innovation Award through the Mayo Clinic Foundation, and the Torrey Coast Foundation.

## Author contributions

This study was conceived of and led by M.L. A.S. designed the model and algorithm with input from M.L., implemented the software, and led data analyses. X.Y. contributed strategies to reduce memory usage and improve computational speed. W.L. provided guidance on handling diverse image formats. M.Y. developed the cell type prediction algorithm. N.S. and L.M. provided the atherosclerosis data, G.X.X. and A.H. provided the melanoma data, B.D. and K.S. provided the kidney data, and T.H.W. provided the gastric cancer data. J.Y. provided advice on data analysis. A.S. and M.L. wrote the manuscript.

## Methods

### Input and image preprocessing

HistoSweep takes a digitized whole-slide histology image *I* ∈ ℝ ^*H*×*W*×3^ as input, where *H* and *W* denote the image height and width in pixels, and the third dimension corresponds to the three RGB color channels (RGB). The image is first rescaled to a user-defined physical pixel size *s* in microns per pixel (default: *s* = 0.5, corresponding to ∼ 20× magnification scanning). The original pixel size from the image metadata, *s*_*raw*,_ is used to compute the appropriate scaling factor *α* where, *α* = *s*_*raw*_ /*s*. Next, padding is applied to the bottom and right edges of the image to allow for complete and consistent patch-wise analysis. Spatial coordinates (e.g., in ST data) are often aligned relative to the top-left origin. Therefore, padding on the right and bottom ensures that the top-left corner of the image, and thus all real tissue coordinates, remains unchanged. The image is then partitioned into non-overlapping square patches (i.e., superpixels) of size *p* × *p* (default: *p* = 16). This results in an effective physical patch size of *sp* × *sp* microns (default: 8µm × 8µm). Therefore, the final rescaled and preprocessed image, defined as *I*′, has dimensions *H*′ and *W*^′,^ such that *p*| *H*′ and *p* | *W*′. Note: We use the terms “patch” and “superpixel” interchangeably throughout the notation.

This flexible patching scheme allows users to tailor HistoSweep to a variety of imaging resolutions and downstream applications. In particular, it supports compatibility with foundation models that operate on fixed-size input/output patches and produce feature embeddings at varying spatial resolutions, while preserving spatial interpretability at the tissue level. Image I/O and resizing are performed using the Python Imaging Library (PIL), while padding and patch reorganization leverage utilities from *skimage* and *einops* for efficient tensor reshaping and manipulation.

### Whole slide patch-level image feature extraction

Following image preprocessing, HistoSweep computes low-level image features for each *p* × *p* patch across the entire tissue section. First, we compute a scalar summary of the color composition for each patch. This provides a compact representation of RGB channel contributions, enabling downstream comparison across heterogeneous tissues. To ensure this metric reflects the most informative aspects of the image, we adopt a channel-adaptive weighting scheme that prioritizes channels with greater global variance, as channels with greater variation are typically more informative for distinguishing tissue structures. As shown in **Supplementary Fig. 9**, global RGB channel variance varies considerably across different tissue types. These observations motivate the use of an adaptive weighting scheme in place of a uniform average. Therefore, we compute the weighted mean intensity *z*_*ij*_ for each patch using the formula introduced in SpaGCN^10^. For each patch location (*i, j*) in the image *I*′, we define the weighted sum *z*_*ij*_ as the following:

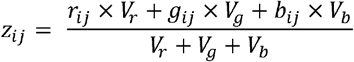

where, *r*_*ij*_, *g*_*ij*_, and *b*_*ij*_ are the mean red, green, and blue intensities for patch (*i,j*)and *V*_*r*_, *V*_*g*_, and *V*_*b*_ are the global variances of the respective RGB channels computed across the entire image.

Furthermore, because the global-variance-weighted intensity formulation does not explicitly capture the within-patch variability, we leverage the standard deviation of pixel intensities across all RGB channels *σ*_*ij*_ to quantify the spatial variation within each patch. For each patch location (*i, j*) we define *σ*_*ij*_ as the following:

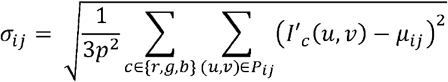

where

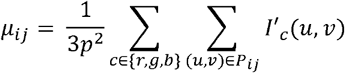

Here *I*′ _*c*_ (*u, v*) denotes the intensity of channel *c* ∈ { *r, g, b* } at pixel location (*u, v*) and *P*_*ij*_ is the set of all pixels in the (*i, j*) -th patch. This local summary statistic serves as a descriptor of color intensity variation within each patch.

To further characterize patch-level variability, we define a ratio metric ℛ_*ij*_ that combines local variation, *σ*_*ij*_, with the global-variance-weighted mean intensity *z*_*ij*._ Specifically, we compute:

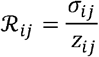

To ensure scale consistency, both *σ*_*ij*_ and *z*_*ij*_ are independently rescaled to [0, 1] via min-max normalization prior to computing the ratio. The ratio metric highlights regions where internal pixel variation is high relative to their overall color intensity, helping to distinguish structured tissue from low-content or uniform regions. As shown in **Supplementary Fig. 10**, this includes histologically informative zones while suppressing background, tissue holes, and fibrotic areas.

### Density-based filtering using joint intensity-variation distribution

To further refine tissue quality and remove non-informative structurally sparse regions, including image artifacts, we apply a density-based filtering strategy based on the joint distribution of the global-weighted intensity *z*_*ij*_ and the local standard deviation *σ*_*ij*_ across all patches. First, both *z*_*ij*_ and *σ*_*ij*_ values are flattened into 1D vectors and used to construct a 2D histogram representing their joint empirical distribution. We then apply a Gaussian filter to smooth the density map and regularize the distribution helping mitigate noise. Each patch is then assigned a density value by binning its corresponding (*z*_*ij*_, *σ*_*ij*_)coordinates into the smoothed histogram. This density reflects how common that patch’s intensity-variation combination is across the entire slide. Finally, patches falling below a given density threshold *t*_*d*_ (default: *t*_*LD*_ = 100) are flagged as low-density (LD). This approach helps identify the set of outlier patches that lie in sparse regions of the joint distribution, typically corresponding to background, debris, poorly stained tissue, or imaging artifacts. We define the set of LD patches as:

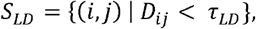

where *D*_*ij*_ is the smoothed joint density estimate assigned to patch (*i, j*) and *τ*_*LD*_ is the density threshold.

### Texture-based refinement of low-density superpixels

To further refine the LD superpixel subset, *S*_*LD*_, and avoid mistakenly discarding informative tissue structures, such as dark nuclei or densely packed cellular regions, we apply a two-stage screening strategy that combines color-based heuristics with texture-based analysis. First, we apply characteristic color signature analysis to automatically exclude superpixels with uninformative or artifactual coloration. Specifically, we discard patches that are (1) overly green (i.e., green is significantly higher than both red and blue: *μ*_*g*_ > *μ*_*r*_ +20 and *μ*_*g*_ > *μ*_*b*_ + 20), (2) near-gray (i.e., low-saturation patches, where all three color channels are similar: ([ *μ*_*r*_, *μ*_*g*_, *μ*_*b*_]) <10) or (3) excessively bright (i.e., each RGB channel mean is high: *μ*_*r*_ > 230 and *μ*_*g*_ > 230 and *μ*_*b*_ >230).

The set of superpixels flagged by this color-based filter is denoted as *S*_*C*_ and will be included in the final LD mask of low-quality superpixels. This partition of the LD set helps remove background debris, tissue folds, or scanning glare before texture-based analysis, enabling more refined and discriminative texture-based clustering. For the following screen, we define the updated LD set as 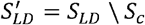 where 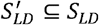.

Next, we perform textural analysis based on gray-tone spatial dependence originally proposed by Haralick et al.^34^. Specifically, HistoSweep performs gray-level co-occurrence matrix (GLCM) texture analysis on patch in the set 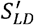. This step aims to increase sensitivity and help avoid false negatives. For each patch in 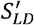, we compute three GLCM-based texture metrics from the corresponding grayscale patch, derived from the RGB histology image. Let *G* be a grayscale image patch of size *p* × *p* in the set *S*_*LD*_, quantized into *L* discrete gray levels (e.g., *L* = 64), where *u,v* refer to gray level bins. The *GLCM* (*u, v*)is a square *L* × *L* matrix defined as:

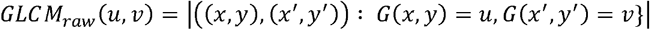

where *x*′, *y*′ is the pixel at a fixed spatial offset from (*x, y*) (e.g. one pixel to the right of (*x, y*)), and | · | denotes the number of times gray levels *u* and *v* co-occur within the patch. Next, to ensure symmetry, we add the transpose of the GLCM to the original matrix, such that 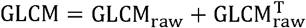. The GLCM is then normalized to form a probability distribution:

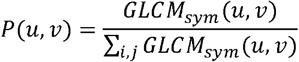

We then calculate the following GLCM-derived texture features for each superpixel in 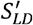.

*Inverse Difference Moment (IDM):*

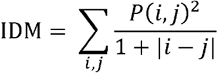

Captures how close GLCM elements are to the diagonal, reflecting local texture smoothness.

*Angular Second Moment (ASM):*

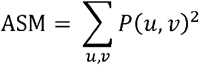

which measures uniformity or pixel pair repetition; high ASM reflects homogenous texture.

*Entropy:*

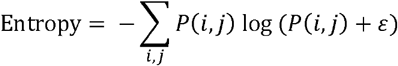

which measures texture complexity or randomness. We add a small constant *ε* to prevent undefined values.

To ensure robustness and interpretability, each metric is min-max normalized to [0,1] across the image. Superpixels are clustered using a Gaussian Mixture Model (GMM) on the three GLCM-derived features (ASM, IDM, and entropy), into four clusters. To automatically label the clusters we define the following composite score as a combination of the GLCM features:

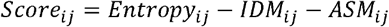

We then use the GMM clustering to partition the set 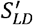 into high-quality superpixels (*S*_*K*_) and low-quality superpixels (*S*_*R*_) selected for removal. The two clusters with the highest average texture scores (*S*_*K*_) are retained, as they exhibit low smoothness (IDM), low homogeneity (ASM), and high entropy, indicating complex, structured regions likely to contain informative tissue. In contrast, the two clusters with the lowest scores (*S*_*R*_) are considered low-quality due to their uniform texture properties. We define the final set of LD superpixels marked for removal as the union of superpixels assigned to low-scoring texture clusters, *S*_*R*_, and the color-screened superpixels, *S*_*C*_. The resulting LD masking set of low-quality superpixels is denoted as:

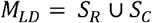

Furthermore, GLCM computation is relatively expensive due to its pairwise intensity co-occurrence nature, especially at scale. By limiting the analysis to this subset, we reduce computational burden while effectively increasing the sensitivity of the mask.

### Ratio-based filtering using Otsu’s threshold

To further refine the superpixel selection, we apply a second stage of filtering based on the global distribution of the normalized ratio metric ℛ. Specifically, we use Otsu’s algorithm^35^ to determine an adaptive data-driven threshold that separates superpixels into groups based on their normalized local variation relative to their weighted mean intensity.

Otsu’s algorithm was originally developed for automatic image thresholding by maximizing the between-class variance in grayscale histograms. In our context, we adapt this method to the distribution of ℛ, seeking to distinguish informative, heterogeneous regions from more homogeneous or uninformative areas. Otsu’s method assumes the input distribution can be approximated by two classes (bimodal) and selects the threshold *t*^*^ that maximizes the between-class variance 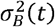 defined as:

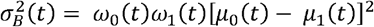

where *ω*_0_ (*t*) and *ω*_1_ (*t*) denote the class probabilities (weights), and *μ*_0_ (*t*) and *μ*_1_ (*t*) are the class means, for the two groups formed by threshold *t*.

To prevent the threshold from being influenced by previously identified low-quality regions, we compute Otsu’s threshold using only superpixels not included in the final low-density mask *M*_*LD*_. Additionally, to mitigate the influence of extreme outliers, we clip the top 10% of values in the distribution of ℛ before applying Otsu’s thresholding.

Let *τ*_*otsu*_ denote the optimal threshold determined by Otsu’s algorithm. Superpixels with *R*_*ij*_ values above *τ*_*otsu*_ are retained, while those below are marked for removal. We define the masking set *M*_*O*_ as:

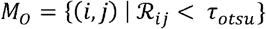

### Connected component analysis and final mask generation

To produce the final binary mask, we first define the set of low-quality superpixels as the union of the LD set *M*_*LD*_ and the Otsu-derived ratio mask *M*_*O*_, such that

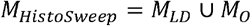

We then construct a binary patch-level mask 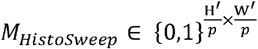, where each superpixel (*i, j*) is marked as 0 if (*i, j*) ∈ *M* _*Histosweep*_, and 1 otherwise.

To further clean the binary mask, we perform connected component analysis in the superpixel space to remove isolated small regions and background debris. Specifically, we apply a morphological filter to remove connected components smaller than an adjustable minimum size (default: 10 superpixels) parameter with 8-connected pixel neighborhood. This step ensures that tiny specks or scanning artifacts are removed.

To generate a pixel-level mask, we unsampled back to the pixel space, each 0 or 1 is expanded into a *p* × *p* block, restoring the full resolution of *H*′ × *W*^′.^

## Data availability

We analyzed the following datasets: **(1)** 10x Genomics whole mouse pup post Xenium H&E image (https://www.10xgenomics.com/datasets/mouse-pup-preview-data-xenium-mouse-tissue-atlassing-panel-1-standard); **(2)** 10x Genomics human colorectal cancer and colon non-diseased Visium HD and H&E image data, IDs: P1 CRC, P2 CRC, P3 NAT, P5 CRC (https://www.10xgenomics.com/products/visium-hd-spatial-gene-expression/dataset-human-crc); **(3)** TCGA-OV — human ovarian serous cystadenocarcinoma H&E images, IDs: TCGA-04-1331-01A-01-BS1, TCGA-25-1320-01Z-00-DX1, TCGA-25-1316-01Z-00-DX, (https://portal.gdc.cancer.gov/projects/TCGA-OV); **(4)** 10x Genomics human lung cancer Visium HD and H&E image data, Experiment 1 (https://www.10xgenomics.com/datasets/visium-hd-cytassist-geneexpression-human-lung-cancer-post-xenium-expt); **(5)** TCGA-LUAD — human lung adenocarcinoma H&E image, ID: TCGA-05-4244-01Z-00-DX1, https://portal.gdc.cancer.gov/projects/TCGA-LUAD; **(6)** human lung atypical adenomatous hyperplasia H&E image and human lung adenocarcinoma 10x visium, 10x xenium, and H&E images generated by Humam Kadara lab, ID: Lung_AAH, Lung_AD (https://doi.org/10.5281/zenodo.17260043); **(7)** human artery atherosclerosis samples post Xenium and CODEX H&E images generated by Lars Maegdefessel lab, ID: Artery_Atherosclerosis (https://doi.org/10.5281/zenodo.17260043); **(8)** TCGA-PRAD — human prostate adenocarcinoma H&E images, IDs: TCGA-YL-A8HO-01Z-00-DX1, TCGA-YL-A8SF-01Z-00-DX1, (https://portal.gdc.cancer.gov/projects/TCGA-PRAD); **(9)** 10x Genomics human liver cancer post Xenium H&E image (https://www.10xgenomics.com/datasets/human-liver-data-xenium-human-multi-tissue-and-cancer-panel-1-standard); **(10)** human kidney non-diseased and T2D post CosMx H&E images generated by Katalin Susztak lab, ID: Kidney_MultiSample_T2D (https://doi.org/10.5281/zenodo.17260043); **(11)** human renal cell carcinoma post mxIF H&E image generated by Alexander Huang lab, ID: Kidney_RCC (https://doi.org/10.5281/zenodo.17260043); **(12)** 10x Genomics human kidney cancer and kidney non-diseased post Xenium H&E images (https://www.10xgenomics.com/datasets/human-kidney-preview-data-xenium-human-multi-tissue-and-cancer-panel-1-standard); **(13)** 10x Genomics human kidney non-diseased Visium HD and H&E image data (https://www.10xgenomics.com/datasets/visium-hd-cytassist-gene-expression-libraries-human-kidney-ffpe); **(14)** TCGA-PAAD — human pancreatic adenocarcinoma H&E image, IDs: TCGA-2J-AAB1-01Z-00-DX1 (https://portal.gdc.cancer.gov/projects/TCGA-PAAD); **(15)** human lymph node from a patient with melanoma post mxIF H&E image generated by Alexander Huang lab, ID: LymphNode_Melanoma (https://doi.org/10.5281/zenodo.17260043); **(16)** 10x Genomics human fresh frozen tonsil with Reactive Follicular Hyperplasia H&E image (https://www.10xgenomics.com/datasets/visium-hd-cytassist-gene-expression-human-tonsil-fresh-frozen-if); **(17)** CCDI human pediatric craniopharyngioma and medulloblastoma brain H&E images, IDs: PBBINC, PBBIHA, (https://ccdi.cancer.gov); **(18)** human MS brain sample CD68 and MOG IHC images^36^, ID: MS330-CAL (https://zenodo.org/records/17261024); **(19)** human gastric cancer Xenium and H&E image data generated by Tae Hyun Hwang lab (https://zenodo.org/records/15164980); **(20)** 10x Genomics mouse non-diseased small intestine H&E image (https://www.10xgenomics.com/datasets/visium-hd-cytassist-gene-expression-libraries-of-mouse-intestine); **(21)** 10x Genomics human breast cancer Xenium and H&E image data, in situ sample 1 replicate 1, in situ sample 2 (https://www.10xgenomics.com/products/xenium-in-situ/preview-dataset-human-breast); **(22)** 10x Genomics fresh frozen human breast cancer tissue (DCIS) H&E image (https://www.10xgenomics.com/datasets/visium-hd-cytassist-gene-expression-human-tonsil-fresh-frozen-if); **(23)** 10x Genomics fresh frozen human invasive ductal carcinoma breast tissue H&E image (https://www.10xgenomics.com/datasets/human-breast-cancer-visium-fresh-frozen-whole-transcriptome-1-standard); **(24)** TCGA-SARC - human sarcoma soft tissue H&E image, ID: TCGA-DX-A6YR-01Z-00-DX1 (https://portal.gdc.cancer.gov/projects/TCGA-SARC); **(25)** 10x Genomics human non-diseased bone marrow tissue H&E image (https://www.10xgenomics.com/datasets/human-bone-and-bone-marrow-data-with-custom-add-on-panel-1-standard); **(26)** TCGA-HNSC - human squamous cell carcinoma mouth tissue H&E image, ID: TCGA-P3-A6T8-01Z-00-DX1 (https://portal.gdc.cancer.gov/projects/TCGA-HNSC); **(27)** mouse mammary gland lymph node H&E slides and 10x Visium ST data collected at two aging time points: 3 months and 18 months^37^, ID: 3M ST1, 3M ST2, 18M ST1, 18M ST2; **(28)** TCGA-THCA - human thyroid carcinoma tissue H&E image, ID: TCGA-ET-A2N1-01Z-00-DX1 (https://portal.gdc.cancer.gov/projects/TCGA-THCA); **(29)** TCGA-BLCA - human bladder urothelial carcinoma tissue H&E image, ID: TCGA-2F-A9KO-01Z-00-DX1 (https://portal.gdc.cancer.gov/projects/TCGA-BLCA). All datasets referenced in this paper have been made publicly available. Details of the datasets analyzed in this paper are described in **Supplementary Table 1**.

## Code availability

The HistoSweep algorithm was implemented in Python and is publicly available on GitHub at https://github.com/amesch441-o1/HistoSweep.

