## Supplementary information for "HistoSweep enables cellular-resolution tissue quality control for gigapixel images in digital pathology and spatial omics"

**Supplementary Table. 1: Datasets analyzed by HistoSweep.**

| **ID** | **Species** | **Tissue Type** | **Data Source** | **Protocol Pre/Post H&E Image** | **H&E image Dimension** |
| --- | --- | --- | --- | --- | --- |
| **S1** | Mouse | Whole mouse pup | (10x Genomics, 2023b),  Pub. date: 08-2023 | (Post) 10x Xenium | 45,056 x 27,392 pixels |
| **S2** | Human | Colon, colorectal cancer | (Oliveira et al., 2024), Sample: p2 CRC | 10x Visium HD | 41,216 x 26,880 pixels |
| **S3** | Human | Colon, colorectal cancer | (Oliveira et al., 2024), Sample: p1 CRC | 10x Visium HD | 39,168 x 32,256 pixels |
| **S4** | Human | Colon, colorectal cancer | (Oliveira et al., 2024), Sample: p5 CRC | 10x Visium HD | 39,936 x 35,328 pixels |
| **S5** | Human | Colon, non-diseased | (Oliveira et al., 2024), Sample: p3 NAT | 10x Visium HD | 44,288 x 23,296 pixels |
| **S6** | Human | Lung, adenocarcinoma cancer | (10x Genomics, 2024c),  Pub. date: 09-2024 | 10x Visium HD, Experiment 1 | 14,080 x 24,064 pixels |
| **S7** | Human | Artery, atherosclerosis | L. Maegdefessel Lab | (Post) 10x Xenium | 20,224 x 14,592 pixels |
| **S8** | Human | Artery, atherosclerosis | L. Maegdefessel Lab | (Post) CODEX | 20,480 x 17,408 pixels |
| **S9** | Human | Multi-sample artery, athero disease prog. | L. Maegdefessel Lab) | (Post) CODEX | 73,728 x 40,704 pixels |
| **S10** | Human | Liver, cancer | (10x Genomics, 2024b), Pub. date: 03-2024 | (Post) 10x Xenium | 8,448 x 18,688 pixels |
| **S11** | Human | Two samples kidney, healthy | K. Susztak lab | (Post) Nanostring CosMx | 40,736 x 34,912 pixels |
| **S12** | Human | Two samples kidney, healthy and type 2 diabetes | K. Susztak lab | (Post) Nanostring CosMx | 42,432 x 37,136 pixels |
| **S13** | Human | Kidney, healthy | (10x Genomics, 2025b), Pub. date: 03-2025 | 10x Visium HD | 24,320 x 17,920 pixels |
| **S14** | Human | Kidney, papillary renal cell carcinoma | (10x Genomics, 2023a), Pub. date: 07-2023 | (Post) 10x Xenium | 4,864 x 15,104 pixels |
| **S15** | Human | Kidney, healthy | (10x Genomics, 2023a), Pub. date: 07-2023 | (Post) 10x Xenium | 6,656 x 18,432 pixels |
| **S16** | Human | Skin, melanoma | A. Huang lab | (Post) mxIF | 22,272 x 27,136 pixels |
| **S17** | Human | Tonsil, reactive follicular hyperplasia | (10x Genomics, 2024d), Pub. date: 09-2024 | 10x Visium HD, fresh frozen | 19,200 x 16,896 pixels |
| **S18** | Human | Gastric, cancer (P1) | T.H. Hwang’s lab | (Post) 10x Xenium | 48,384 x 19,712 pixels |
| **S19** | Human | Gastric, normal (P1) | T.H. Hwang’s lab | (Post) 10x Xenium | 48,384 x 24,064 pixels |
| **S20** | Human | Gastric, normal (P2) | T.H. Hwang’s lab | (Post) 10x Xenium | 33,280 x 23,808 pixels |
| **S21** | Mouse | Small intestine, healthy | (10x Genomics, 2024e), Pub. date: 03-2024 | 10x Visium HD | 13,056 x 12,032 pixels |
| **S22** | Human | Breast, cancer | (Janesick et al., 2023), Replicate 1 | (Post) 10x Xenium | 20,224 x 15,104 pixels |
| **S23** | Human | Breast, cancer | (10x Genomics, 2025a), Pub. date: 02-2025 | 10x Visium HD, fresh frozen | 17664 x 16128 pixels |
| **S24** | Human | Breast, invasive ductal carcinoma | (10x Genomics, 2022),  Pub. date: 07-2022 | 10x Visium, fresh frozen | 20,992 x 14,080 pixels |
| **S25** | Human | Bone marrow, nondiseased | (10x Genomics, 2024a), Pub. date: 04-2024 | (Post) 10x Xenium | 20,000 x 7,536 pixels |

**Supplementary Fig. 1: Ratio norm** **distributions and Otsu’s thresholds across different samples.** Histograms of normalized ratio values across samples from four spatial omics platforms. Each panel shows the distribution of ratio norm values with the corresponding Otsu-derived threshold (red dashed line) used for downstream filtering. Samples include **a,** VisiumHD (S2: colorectal cancer), **b,** post CosMx (S12: kidney normal + T2D), **c,** post Xenium (S12: gastric normal), and **d,** post CODEX (S8: artery atherosclerosis).

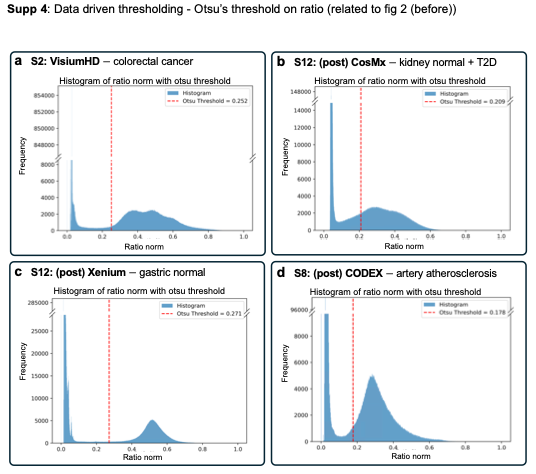

**Supplementary Fig. 2: Application of HistoSweep on extensive artery dataset.** HistoSweep processing of a massive post-CODEX H&E image. Left: H&E-stained whole-slide image spanning 15 artery samples. Right: Corresponding HistoSweep output, generated in minutes on a CPU. The full image exceeds 3 billion pixels (73,728 × 40,704) at 0.5 µm/pixel resolution.

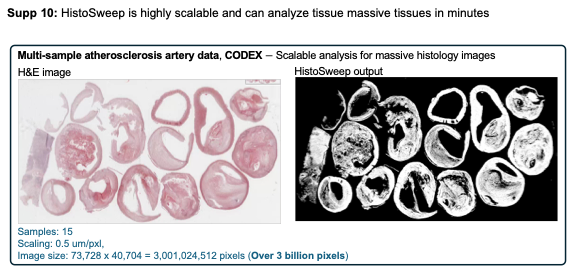

**Supplementary Fig. 3: HistoSweep mask for post-Xenium whole mouse pup H&E image.** Top panel shows the original H&E image, and bottom panel shows the HistoSweep generated tissue mask.

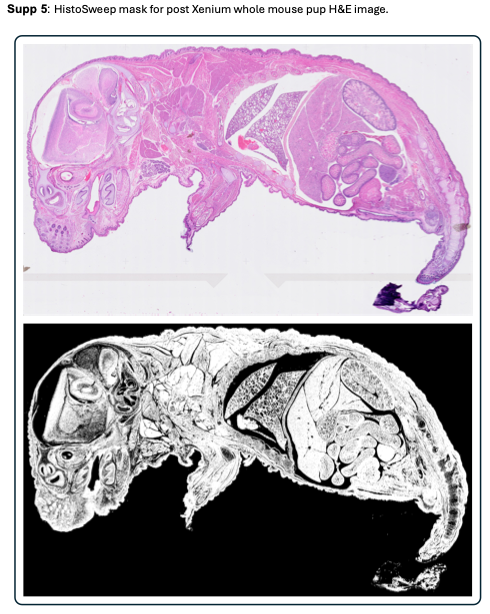

**Supplementary Fig. 4: Detecting signal leakage in Visium HD colorectal cancer data**. Analysis using only “bins under tissue” determined by Visium HD software. **a,** H&E images, Visium HD total UMI counts (8 µm bins), and HistoSweep removal and retention maps. Insets show magnified regions highlighting local histology and corresponding transcript counts. **b,** Proportion of bins retained and removed by HistoSweep (left), violin plot showing total UMI by HistoSweep status (right), and density distributions of total UMI counts across retained and removed bins (bottom).

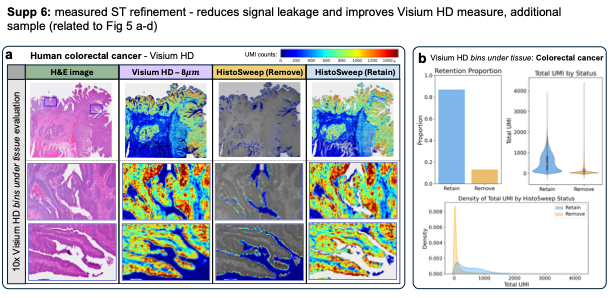

**Supplementary Fig. 5: Evaluation of HistoSweep specificity and UMI distributions across all Visium HD bins. a, c, e**, Comparison of Visium HD tissue detection masks with HistoSweep filtering results for healthy kidney (a), lung cancer (c), and colorectal cancer (e) samples. Regions not detected as tissue by Visium HD are labeled as “Not tissue” (serving as true negatives). **b, d, f,** Density plots and violin plots of total UMI counts by HistoSweep status across all bins in each sample.

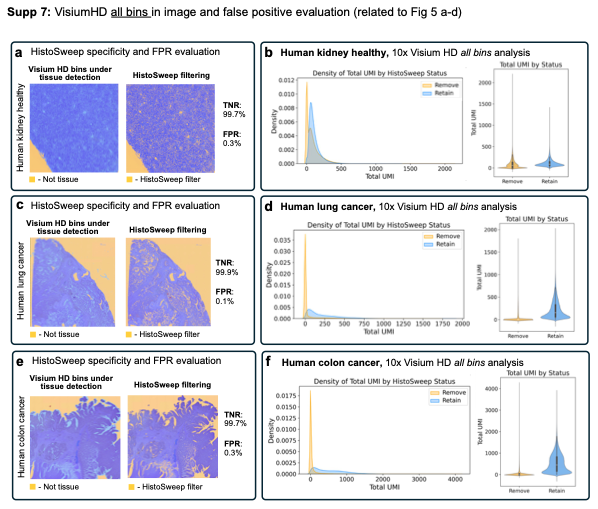

**Supplementary Fig. 6: Spatial metric visualizations and UMI comparisons for HistoSweep groups in Xenium gastric samples.** **a,** H&E images with corresponding scatterplots of RGB standard deviation versus weighted mean colored by superpixel density, ratio statistic, and total UMI plots for one tumor and two normal gastric samples. **b,** Violin plots showing total UMI distribution in retained vs. removed regions within Xenium-measured tissue boundaries. **c,** Violin plots showing total UMI distribution in retained vs. removed regions across the entire histology image.

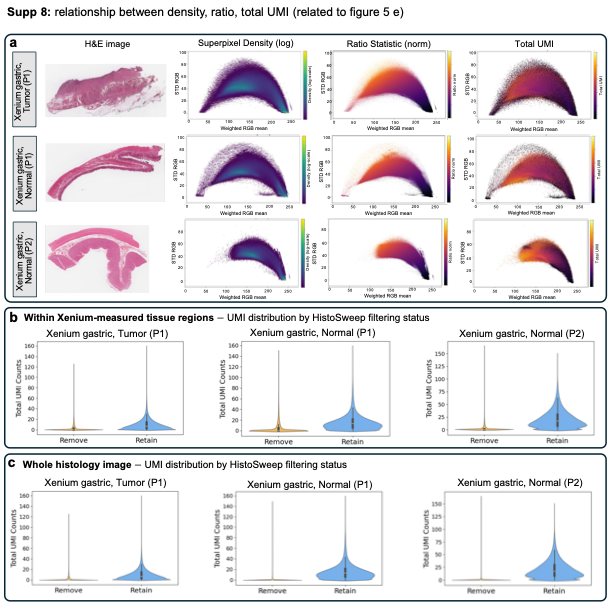

**Supplementary Fig. 7: Evaluation of high-UMI filtering and alignment quality in Xenium gastric normal samples.** **a, b, d,** Total UMI vs. ratio statistic density plots for a gastric cancer sample and two normal gastric samples (right), with superpixels in the low-ratio, high-UMI region highlighted and spatially mapped (left). HURR = 6.33%, 3.12%, and 0.66%, respectively. **c, e,** H&E images with overlays of filtered high-UMI superpixels (red), HistoSweep mask application, and aligned Xenium transcript locations showing differences in alignment quality between samples.

**Supplementary Fig. 8: Extension of HistoSweep to IHC stained histology images in post-mortem human brain tissue with multiple sclerosis. a,** IHC-stained section for CD68 highlighting activated microglia/macrophages. **b,** Adjacent section stained for MOG marking myelinated regions. Right panels show HistoSweep detection using the current low-density pipeline without adaptive thresholding on the ratio. Original IHC data were reported in Schroeder et al. 2025 Nature Methods, PMID: 40060412.

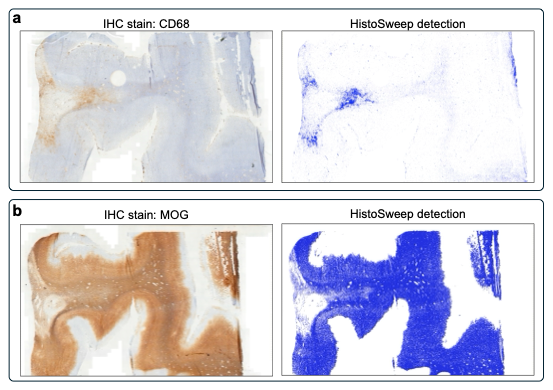

**Supplementary Fig. 9: Global RGB channel variance across different samples.**

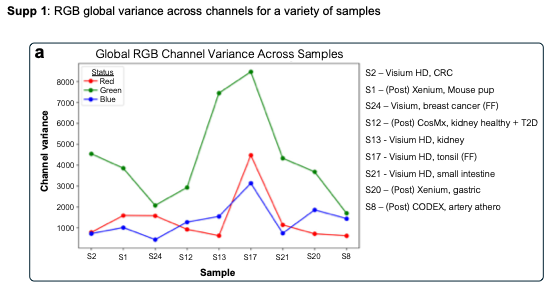

**Supplementary Fig. 10: Visualization of H&E, RGB channel intensity, and calculated ratio metric for each sample.**

H&E	Red channel	Green channel	Blue channel	Ratio metric

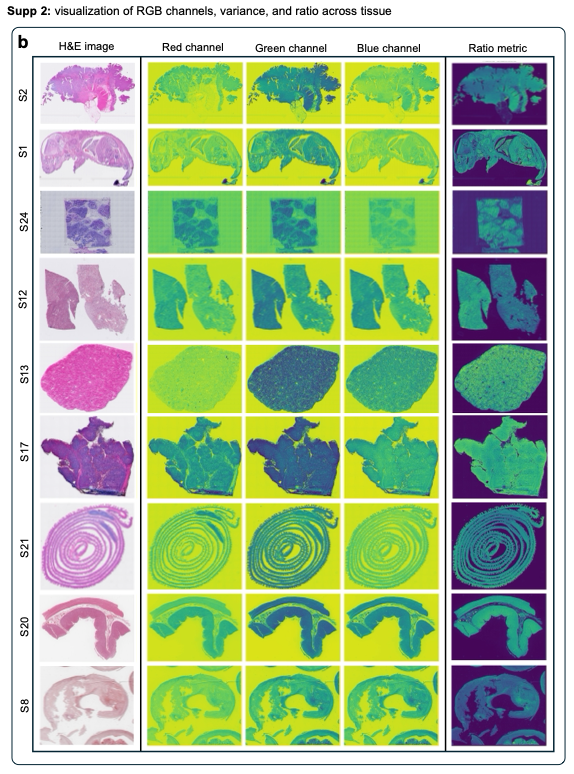
